# Mineralized coleoid cranial cartilage from the Late Triassic Polzberg *Konservat-Lagerstätte* (Austria)

**DOI:** 10.1101/2022.02.14.480445

**Authors:** Petra Lukeneder, Alexander Lukeneder

## Abstract

Although hyaline cartilage is widely distributed in various invertebrate groups such as sabellid polychaetes, molluscs (cephalopods, gastropods) and a chelicerate arthropod group (horseshoe crabs), the enigmatic relationship and distribution of cartilage in taxonomic groups remains to be explained. It can be interpreted as a convergent trait in animal evolution and thus does not seem to be a vertebrate invention. Due to the poor fossil record of cartilaginous structures, occurrences of mineralized fossil cartilages are important for evolutionary biology and paleontology. Although the biochemical composition of recent cephalopod cartilage differs from vertebrate cartilage, histologically the cartilages of these animal groups resemble one another remarkably. In this study we present fossil material from the late Triassic Polzberg *Konservat-Lagerstätte* near Lunz am See (Lower Austria, Northern Calcareous Alps). A rich Carnian fauna is preserved here, whereby a morphogroup (often associated with coleoid remains) of black, amorphous appearing fossils still remained undetermined. These multi-elemental, mirroring fossils show remarkable similarities to recent cephalopod cartilage. We examined the conspicuous micro- and ultrastructure of these enigmatic fossils by thin-sectioning and Scanning Electron Microscopy (SEM). The geochemical composition analyzed by Microprobe and Energy Dispersive X-ray Spectroscopy (SEM-EDX) revealed carbonisation as the taphonomic pathway for this fossil group. 3D preservation of this otherwise degradable soft tissue can be explained by the mineralization processes that may take place in particular environments. Fifty-nine specimens from the Polzberg locality and five specimens from Cave del Predil (formerly Raibl, Julian Alps, Italy) underwent extensive studies. The detected grooves are interpreted to be muscular attachment areas, and the preserved branched system of canaliculi is comparable to a channel system that is also present in recent coleoid cartilage. The new findings on these long-known enigmatic structures strongly point to the preservation of cranial cartilage belonging to the belemnoid *Phragmoteuthis bisinuata*.

## Introduction

Hyaline cartilage as connective tissue with its supporting, skeletal function has a long evolutionary-biological history and was developed in many different animal lineages such as polychaetes, echinoderms and molluscs [1-5]. Within the molluscs, hyaline cartilage is reported as a radula-supporting structure in predatory gastropods such as *Busycon canaliculatum*^5^ and various cephalopods [2-7]. The fossil record of cartilage and cartilaginous structures is sparse.

Cranial cartilage of an early Jurassic cephalopod *Loliginites* (*Geoteuthis*) *zitteli* (= *Loligosepia aalensis*) [8] was reported from the Posidonia shale of Schömberg (Germany) [9], later re-described [10]. Jurassic coleoid specimens from Solnhofen limestone comprise preserved cephalic cartilage including statocysts and other soft tissues [11]. From the Polzberg *Konservat-Lagerstätte*, the preservation of coleoid soft tissues and probable mandibles of *Phragmoteuthis* specimens, as well as of Trachyceratid soft tissues, has already been addressed [12,13].

This study examines historical fossil material and new findings from the Polzberg locality and Cave del Predil. Earlier research suggested that the enigmatic black structures from Cave del Predil were halves of coleoid jaws [14,15], belonging to *Acanthoteuthis bisinuata*. The presence of cartilaginous rings on the slabs was also proposed [14]. This theory was later adopted by numerous authors [Doguzhaeva, pers. com. 2012; 16,17] but this contribution is the first to provide concrete evidence.

### The Evolution of invertebrate cartilage

Hyaline cartilage with its translucent appearance and the cartilage cells embedded in an extracellular, hydrophilic matrix occurs not only in vertebrates, but is also widely distributed in various lophotrochozoan groups [1] such as in some gastropod buccal masses, cephalopods, and sabellid polychaetes [1,2,3,5,6,18-20]. Outstanding is the occurrence of real cartilage in horseshoe crabs. The assumption is therefore that this tissue is not a vertebrate invention [3,6,20] but evolved convergently more than once. Although invertebrate cartilage differs biochemically from vertebrate cartilage^5^, cephalopod and vertebrate cartilage share similar morphological features on the histological level [2,5,19,21,22]. Cephalopod cartilage even shares similarities to vertebrate bone [23]. The initial cartilaginous structures visible at the embryonic stage are the funnel cartilage and the cartilage of the pallial complex within the cephalopod brain [6]. Even in hatchlings, cranial cartilage is visible as a thin layer of collagen in hatchlings of certain cephalopod groups such as *Loligo pealeii* [6]. The recent decapods (“ten-armed coleoids“), to which *Sepia* spp. also belong, are close relatives of belemnite-resembling cephalopods with a weakly developed rostrum [24]. In contrast to other recent cephalopods only in the sepiid coleoids and the deep sea cephalopod group of *Spirula spirula* still have kept their mineralized phragmocone [25]. The predatory cephalopod mode of life requires an early development of additional supporting and protective functions of the ocular cartilages and an oculomotoric sensory system [6,26]. Several cartilage types are associated with the eye [1]: cranial cartilage, scleral cartilage and the equatorial ring (“iris-cartilage”) [4,19]. Ocular cartilage is present in several cephalopod groups such as *Nautilus, Octopus, Elodene, Sepia* and Spirula [5]. The scleral cartilage and the equatorial ring contain less matrix than the cephalic cartilage, which is also a less cellular type [4]. Conversely, eye-associated cartilage seems to be more cellular (containing less matrix) than cranial cartilage [19].

### The structure and function of cranial cartilage in recent squid

Thirteen cartilaginous structures are present in adult specimens of *Sepia officinalis*, most function as attachment points for muscular structures and are therefore important for the locomotory system [6]. Magnetic resonance imaging data of different cephalopods provides insights into the spectrum of cephalopod cranial cartilage [27]. The supporting function of cranial cartilage (= cephalic cartilage = head cartilage [1]) protects the squid’s brain from external forces [4]. This structure is incomplete [5], featuring orbital depressions [4,5]. Extensive cephalic cartilage is present in several cephalopod groups such as *Octopus vulgaris, S. officinalis* (European cuttlefish) and *Loligo pealii* [4]. Optical ganglia of the nervous system and the eyes are housed in ocular cartilage, which can be interpreted as lateral parts of the cephalic cartilage [6]. These cartilaginous parts become fused in late developmental stages by bridging cartilage and statocysts, which contain the statoliths [6]. Extensive research [1,3,5,6,20,21,28,29] shows that the cartilaginous ultrastructure of several cephalopod groups histologically resembles vertebrate cartilage. The funnel cartilage of *I. illecebrosus* strongly resembles vertebrate cartilage [29]. Cartilaginous structures such as the funnel cartilage, but also the cephalic cartilage in *S. officinalis*, are surrounded by a distinct, thin layer, the perichondrium [3]. The occurence of cartilage-lining cells indicates cell growth from the central cartilage part [3]. The presence of a channel system is one similiarity between coleoid and vertebrate cartilage [6]. This system isoften present in the extracellular matrix of cephalopod cranial cartilage as a passage for blood vessels [1,4]. Interestingly, the coleoid chondrocytes (cartilage cells) extend into long processes [6,30,31], a feature typical of vertebrate osteocytes but not for vertebrate chondrocytes. Furthermore, the morphology of cephalopod chondrocytes seems to change relative to their particular position within the cartilage[23].

Several authors reviewed the biochemical composition of recent cephalopod cartilage [4,6]. The extracellular matrix of cephalopod cartilage consists of fibrous proteins in a hydrophilic ground substance [6]. The extracellular matrix of *Sepia* contains different types of collagen [30]. The staining of this matrix in *Illex illecebrosus* is unevenly distributed [19]

Various authors examined the mineralization processes of invertebrate cartilage in media metastable to hydroxylapatite at 37°C [32] and references therein. The histology and mineralization of the cephalic cartilage of *L. pealii* was figured in [4]. A peripheral thickening in some squids was determined in reconstructions of the microanatomy and histology of the central nervous systems and of the eyes in coleoid hatchlings [33].

### The fossil record of cephalopod cartilage

Fossilisation includes processes in which organic components are replaced by inorganic minerals. Although the fossil record of cartilage ranges back to the Paleozoic [34], it is poor due to the rapid degradation of soft tissues. Cenozoic fossil cartilage, where the exceptional preservation state was linked to a surrounding phosphate-rich environment, can be examined using immunohistochemical and biochemical methods [35]. Calcified cartilage of the Campanian (Upper Cretaceous) *Hypacrosaurus stebingeri* was the subject of more recent investigations [36,37]. Early historical investigations assumed coleoid cephalic cartilage and its muscular elements in a specimen of “*Loliginites zitteli*“ (= *Loligosepia aalensis* [8]) and described this in detail [9,10,38] figured cephalic cartilage from different fossil coleoids, among them *Plesioteuthis prisca* and *Loliginites zitteli*. An Upper Carboniferous cranial cartilage capsule was reported from Oklahoma [39]. Important findings on the coleoid *Dorateuthis syriaca* from *Konservat-Lagerstätten* in Lebanon enabled reconstructing the cephalic cartilage of this taxon [40]. A detailed compilation of fossilised cranial cartilage findings within the coleoid genera was more recently published [41]. Accordingly, indications for fossilised head cartilage were found in:

1. the phragmocone-bearing coleoid genera *Acanthoteuthis* and possibly *Phragmoteuthis*,
2. the gladius-bearing Prototeuthina *Plesioteuthis, Dorateuthis, Boreopeltis, Rhomboteuthis*,
3. the Loligosepiina *Vampyronassa, Proteroctopus, Massigophora*, and
4. the Teudopseina *Trachyteuthis, Glyphiteuthis, Rachiteuthis* and *Muenstella*.

More recently, cranial cartilage with statocyst remains in specimens of a Jurassic *Acanthoteuthis* from Solnhofen (Germany) were examined [11]. Fin cartilages of the middle Olenekian (Lower Triassic) coleoid *Idahoteuthis parisiana* from Idaho (USA) revealed a canalicular structure [42]. UV-light and light of visible wavelengths was useful in studying the cephalic cartilage of carboniferous coleoids [43], where the orange color of parts of the head cartilage indicated phosphatised soft tissue.

### The Polzberg *Konservat-Lagerstätte*

The paleontological site Polzberg has been known for almost 150 years and was reported synonymously under various names such as Unter Polzberg, Pölzberg, Polzberg-Graben, Schindelberggraben and Polzberg [16,17,44] and references therein. In early collections, material from deposits of the Reingraben Shales was not separated from the Lunz Formation, and analogous specimens from the Polzberg locality were often designated as Lunz locality [17, 44]. The fossil site is located 4.5 km north-east of Lunz am See, on the western slope of mount Schindelberg (= mount Schindlberg, 1066 m), within the Reifling basin and belongs to the Bajuvaric Lunz Nappe System of the Northern Calcareous Alps (Fig. 1). It is accessible from the north over Erlauftal street 25, then Zellerrain street 71 or from the south via Mariazell over the Zellerrain street 71. The exact GPS position is N 47°53’5.90’’ and E 15° 4’27.70’’ (see also [17]; 712 m above sea level).

**Figure 1.**
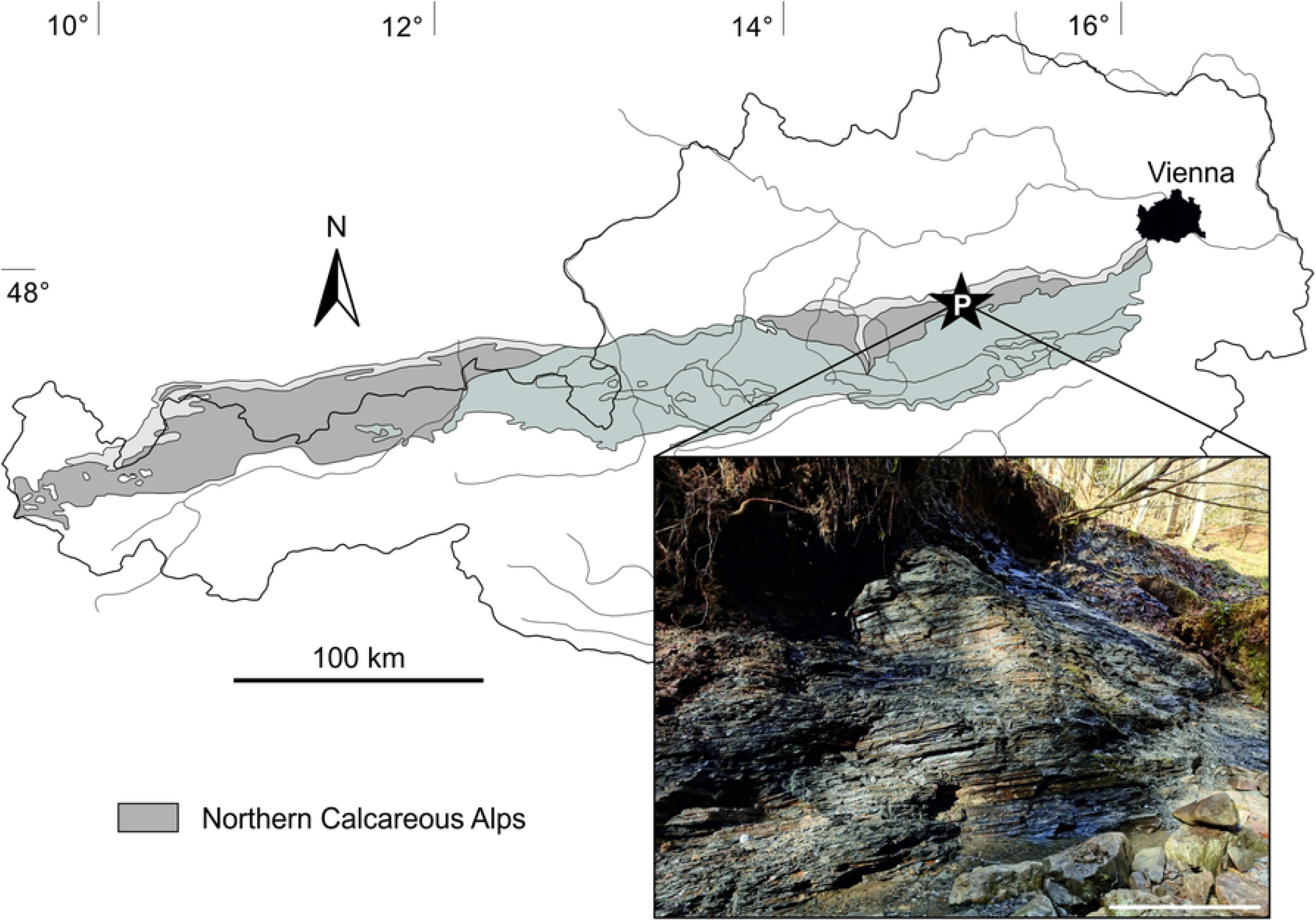
Austrian map with Northern Calcareous Alps marked. Black star: position of the *Konservat-Lagerstätte* Polzberg (= Schindelberggraben ravine, Polzberg locality) near Lunz am See with positions of new outcrops and historical adits. Prepared by PL and AL using CorelDRAW X7; www.coreldraw.com.

The IRIS system (Interaktives Rohstoffinformations System) hints to an early historical adit (No. 071/3008a) for the production of black coal, active in the first half of the 19^th^ century, which was confirmed by the literature [45]. The outcrop Polzberg locality (Fig. 2A) was first mentioned by [45]. Further historical adits for the recovery of fossil specimens were dug in 1885 (Geological Survey of Austria, GBA) and in 1909 (Natural History Museum Vienna, NHMW) by Joseph Haberfelner [17] and references therein at approx. GPS 47°53’6.20”N, 15° 4’28.30”E (710 m a.s.l.; Fig. 1). In early research, the deposits outcropping at Polzberg were assigned to the Wengen shales [45], bearing fossils such as *Ammonites aon*, the double-valved crustacean *Eusteria* sp. and specimens that belong to the highly variable taxon *Halobia minuta*. Coleoid specimens of *Acanthoteuthis bisinuata*, fish remains of *Belonorhynchus striolatus* and plants assigned to *Voltzia haueri* were also reported [45]. Dark-grey to black, foliated clays and marls of the Reingraben Shales (= *Halobienschiefer*) intercalated by several limestone beds crop out at the Polzberg locality. Exceptionally preserved fossils are distributed within these sediments, pointing to an fully marine origin with sporadic influx of freshwater [46]. The older Reingraben Shales are thought to be a deep marine environment with nektonic (actively free-swimming) faunal elements [46]. For the Polzberg *Konservat-Lagerstätte*, dysoxic to anoxic conditions were proposed [44], while other authors favored a shallow marine, basinal environment [12]. The enigmatic fossils from Cave del Predil were described and figured in early works and assigned to the coleoid *Acantotuethis bisinuata* [14]. Historical excavations by the GBA and the NHMW, as well as recent findings by fossil collectors (2012-2014) and scientific field work in 2021, yielded similar specimens (Fig. 2B) from Polzberg.

**Figure 2.**
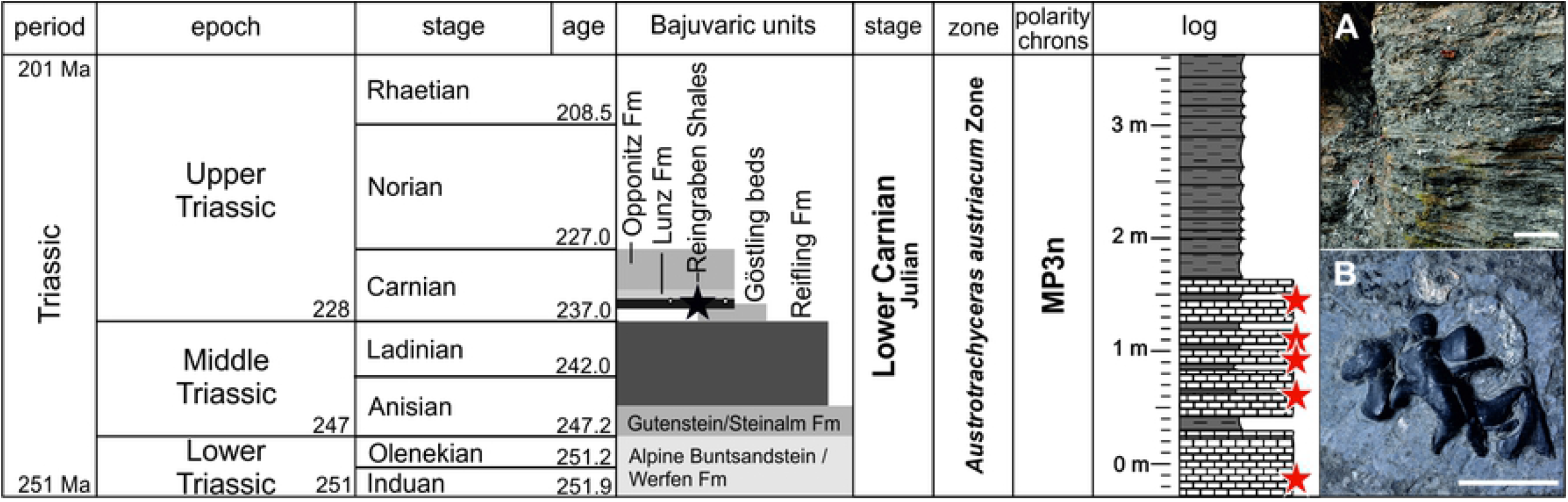
Stratigraphic chart with Bajuvaric Formations and lithological overview log of Polzberg locality; calcareous deposits occur only in the older/lower part of the section. **(A)** Vertical section (middle part) of Polzberg locality. Scale bar 20 cm. **(B)** Enigmatic carbonised structure from Polzberg locality, NHMW 2021/0001/0002. Layers with occurring carbonised elements marked by red asterisks. Scale bar 1 cm. Prepared by PL and AL using CorelDRAW X7; www.coreldraw.com.

## Material and methods

### Material and institutional abbreviations

NHMW, Natural History Museum Vienna, GBA Geological Survey of Austria. Sixty-four specimens of “cartilagenious“ structures were available for analyses. A full list of examined specimens is provided in S1 Tab. All specimens were drawn digitally and measured. One specimen was used for thin-sectioning (NHMW 2012/0117/0024), 13 specimens were analyzed by serial sectioning. 59 specimens stem from the Carnian Reingraben shales at the Polzberg locality (= Polzberggraben; historical and new findings). Bed-by-bed collected specimens found by the authors from distict levels: Po -50–0 cm, Po 60–80 cm, Po 80–100 cm, Po 100–120 cm, Po 140–160 cm (Fig. 2). Five fossils come from Cave del Predil (northern Italy). The coleoids from the Polzberg *Konservat-Lagerstätte* are compressed and flattened, while the investigated black, enigmatic structures are not. The analyzed material is stored in the paleontological collection of the NHMW and the GBA. The fossil structures were analyzed with a variety of analytical tools as follows:

### Digital imaging and image processing

Macro-photography of all fossil specimens was done with a Nikon D 5200 SLR, lens Micro SX SWM MICRO 1:1 Ø52 Nikon AF-S, Digital Camera, in combination with the freeware graphic tool digiCamControl Version V.2.1.2.0 at the NHMW. High-resolution digital micro-photography was done using a Discovery.V20 Stereo Zeiss microscope, processed with the software AxioVision SE64 Rel. 4.9 imaging system at the NHMW.

### Scanning Electron Microscopy (SEM)

SEM images of the surface, as well as of the internal structures, of specimen NHMW 2012/0117/0024 were taken using the Quanta™ 250 FEG from FEI (with a Shottky field emission source FEG-ESEM) from the Department of Material Sciences and Process Engineering (MAP) at the University of Natural Resources and Life Sciences, Vienna. The electron microscope was equipped with an Everhardt Thornley SED-Detector, in low-vacuum settings with 15kV accelerating voltage. The specimen was therefore not gold-sputtered. Overview images were taken with a JEOL “Hyperprobe” JXA 8530-F field-emission electron microprobe (FE-EPMA) in combination with an online JEOL quantitative ZAF-correction program at the Central Research Laboratories of the NHMW. The sample was coated for EDS analyses with an 8 nm carbon film. An accelerating voltage of 15 and 5 keV, a beam current of 5 nA, and fully focused electron beam (beam diameter of approx. 70–80 nm) were used. The Count Rate was 1055.00 CPS. EDS studies of carbonised structures and sediment were performed with an FEI Inspect-S scanning electron microscope with an EDAX Apollo XV SDD EDS detector at 15, 10 and 5 keV acceleration voltage. Spectra were acquired for 30-90 s to obtain a good signal to noise ratio, and intensities were corrected with the ZAF algorithm. One piece of the specimen was sputter-coated with gold and scanned with high-voltage. EDS-SEM results can be taken from S3 Fig. for carbonaceous material and from S4 Fig. for calcitic fillings.

### Micro Computertomography (Micro-CT)

Thirteen selected specimens were serially sectioned at the Core Facility for Micro-Computed Tomography (Vienna Micro-CT Lab), University of Vienna, Austria, using the custom-built VISCOM X8060 NDT (Germany) µ-CT scanner with different scan parameters, which delivers a stack of images with isometric voxel sizes. These are then combined, resulting in a 3D volume. Scan parameters for each specimen can be taken from S4 Tab.

The specimens were segmented using the software Avizo Amira 2020 (Thermo Fisher Scientific) and virtually 3D-reconstructed in Drishti 2.7 [47].

### Mineralogical and geochemical studies

The composition of one specimen was analyzed by Raman spectroscopy at the Department of Mineralogy, University of Vienna. Raman spectroscopy was used for the in-depth characterisation of the carbon phase of the fossil remains. The Raman spectra were obtained by means of a Horiba LabRAM HR Evolution system equipped with an Olympus BX-series optical microscope and Si-based charge-coupled, Peltier-cooled, device detector. Spectra were excited with 532 nm emission of a frequency-doubled Nd-YAG laser (calcite, 12 mW at the sample; graphitic carbon, 0.012 mW). A 50x lens (numerical aperture 0.55) was used to focus the light onto the surface of the sample. The light to be analyzed was dispersed with 1800 grooves per mm diffraction grating. Raman spectroscopy for the specimen NHMW 2021/0016/0397 (S5 Fig.) revealed strongly disordered (i.e., sp^2^ hybrid bonded) carbon as the material of the fossil structures. They were measured at low energies to avoid measuring the burning spot. The fossils were measured in comparison to reference spectra of natural graphite from a uranium mineralization in Saskatchewan, Canada (for sample description see [48, 49]).

Elemental mapping with the Microprobe JEOL Hyperprobe JXA-8530F field emission electron microprobe (EMS) in combination with the online JEOL quantitative ZAF-correction program was conducted at the Central Research Laboratories of the NHMW on specimen NHMW 2012/0117/0024 (S6 Data).

### Measurements and statistics

Where possible, specimens were measured by using a digital caliper (S7 Fig., S8 Tab.). Further detailed measurements models were carried out on original Micro CT data using the software Dragonfly Workstation Version 2021.1 and Avizo Amira V. 5.4.0. Measurements for comparison with recent *Sepia officinalis* were done on a dataset from Ziegler et al. [27]. Diameters of the fossil channel system were measured on original data of a specimen with a well-preserved canalicular system (NHMW 2012/0117/0028, at a resolution of 15 µm). Statistics was done with Microsoft Excel 2010.

### Thin-sectioning

Thin-sections from one specimen (NHMW 2012/0117/0024) were prepared at the laboratory of the Natural History Museum Vienna, Austria. The specimen was embedded in Araldite epoxy resin, then sectioned and mounted on slides for microscopy. The sections were polished with aluminium oxide and silicon carbide powders to a thickness of about 25 μm.

### Sections of recent squids

Sections of three dead squid *Loligo vulgaris* and two *Sepia officinalis* yielded deep insights into the morphology of cephalopod cartilage and enabled actuopaleontological comparisons. Anatomical sections of the coleoids were produced at the Department of Palaeontology, University of Vienna, Austria, and the cranial cartilages were isolated. Coleoids and cartilage were measured and photographed.

## Results

Within the Polzberg section, the investigated fossils were only found in the lower part of the section and thus seem to be limited to the more calcareous part (layers -50 to 0 cm, 60 to 80 cm, 80 to 100 cm, 100 to 120 cm and 140 to 160 cm) (Fig. 2, S1 Tab.).

Two distinct, morphologically different types of black, amorphous, carbonised masses (Fig. 3, S2 Fig., S5 Fig.) with a canal system (= channel system, canalicular system) were recorded – Type A (45 specimens) and Type B fossils (10 specimens). Type A fossils sometimes occur with a “wing-like” structure (9 specimens). Isolated “wing-like” elements only occur in four specimens. Seven fossils come from a private collection, 14 were new findings from recent excavations, 32 belong to historical material from the Natural History Museum Vienna (NHMW, Austria). The five examined specimens from the Geological Survey Austria (GBA) come from Cave del Predil. 14 specimens (13 from Polzberg locality, one from Cave del Predil) featured associated, mainly *in-situ*-preserved coleoid microhooks, eight specimens exhibited a coleoid proostracum. We found four enigmatic fossils with proostracum and hooks. All specimens show a surrounding calcitic sea/layer (approx. 20 μm thick) with a preserved, calcite-filled channel system. Both types mainly appear in the lower, calcareous part of the section (S1 Tab.). S1 Tab. further summarizes the analytical methods used on the examined specimens from both localities. Measurements on the particular elements are listed in S8 Tab.

**Figure 3.**
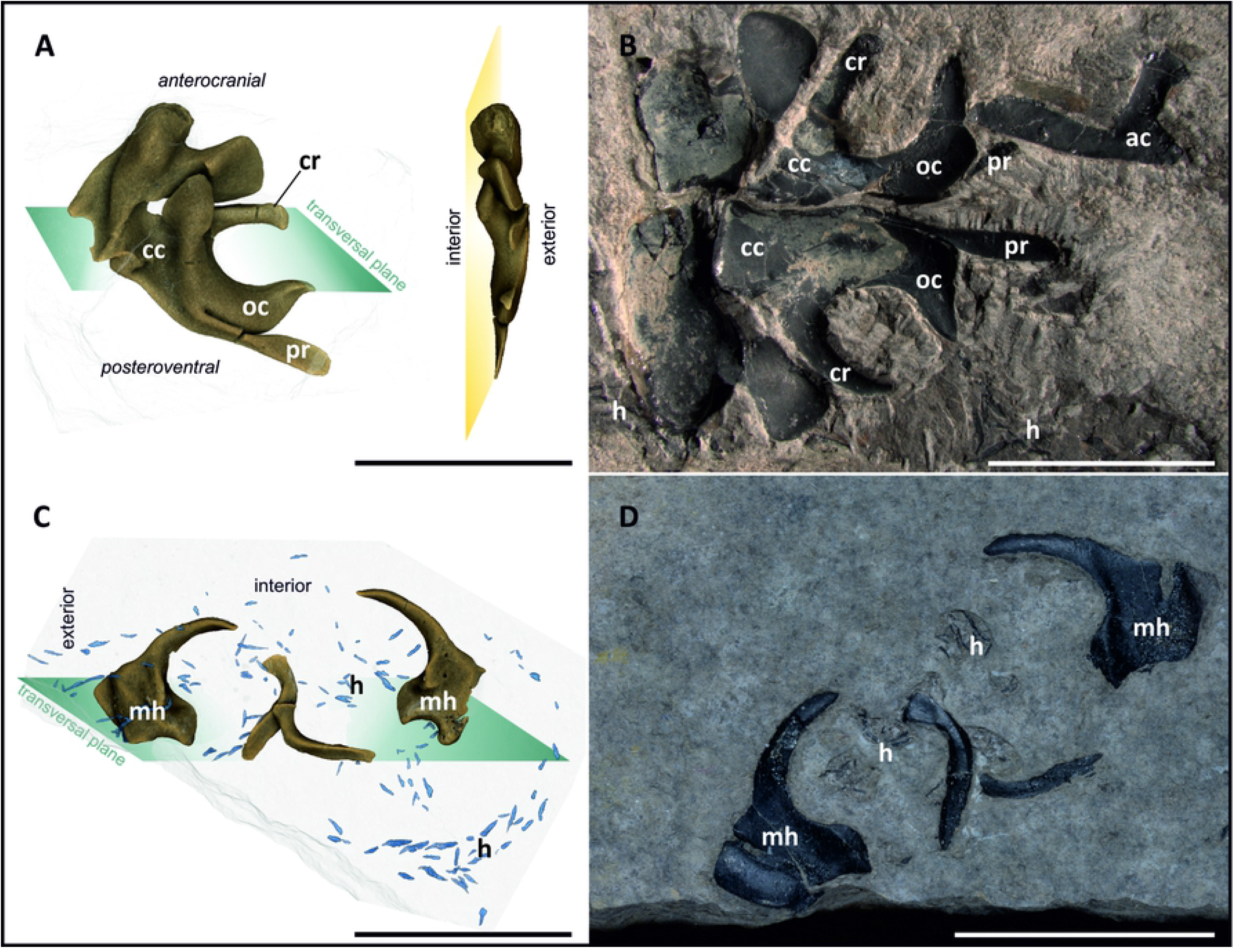
Specimens of Type A and Type B fossils from the Polzberg *Konservat-Lagerstätte*. **(A)** Suggested orientation for Type A fossils showing characteristic, curved C structure with prolonged processus. **(B)** Most entire specimen showing mirroring, semi-*in-situ* configuration of the structures with likewise *in-situ* preserved coleoid hooks, starting in the area of the long dorsal processus above the C structure. **(C)** Suggested orientation for Type B fossils showing a curved hook-like structure. **(D)** Mega-hook-like structures associated with coleoid hooks, NHMW 2021/0124/0003a, b. All scale bars 1 cm. ac arm cartilage, cc cephalic cartilage, cr carrier, oc ocular cartilage, h coleoid hooks, mh mega-hook structures, pr processus. Prepared by PL using CorelDRAW X7; www.coreldraw.com.

### Type A Fossil

Type A fossils (Fig. 3A, B and 4A, B). Black, shiny, bilateral structure, maximum fossil lengths (diameter from most proximal point at processus to lowermost part of supposed “statocysts“) size from 5 mm (single element) up to 24 mm (double elements); particular elements of the specimens such as carrier“ (Fig. 3A, B) appear loosely connected; calculated volumes range from 31.82–253.95 mm^3^ depending mainly on completeness of the available structure.

**Figure 4.**
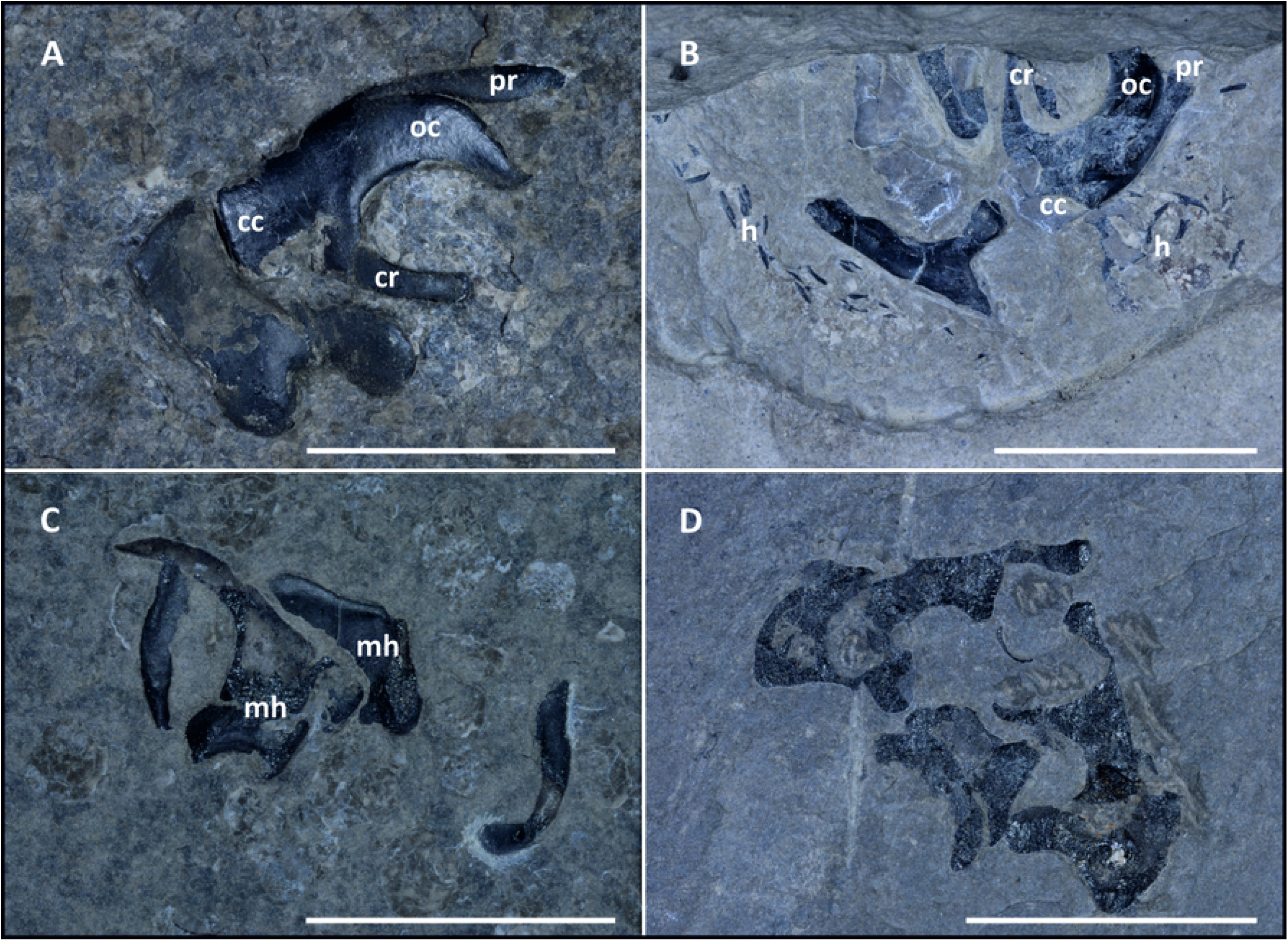
Different views of Type A (possible cranial cartilage) and B specimens (megahooks). **(A)** Standard »lateral« view of Type A fossils (NHMW 2012/0117/0001) with groove, characteristic processus on prominent C structure. **(B)** Supposed »cranial« view of Type A specimen (NHMW 2012/0117/0025) with associated wing-elements and *in-situ* preserved hooks. **(C)** Two megahooks with a triangular base and two additional elongated elements (NHMW 2021/0124/0002). **(D)** Historical specimen from the Geological Survey Austria (GBA 2006/011/0041) from the Rinngraben locality near Cave del Predil (Raibl, Italy). These specimens show a thicker and more prominent upper part of the C structure. cc cephalic cartilage, cr carrier, h coleoid hooks, mh mega-hook structures, oc ocular cartilage, pr processus. All scale bars 1 cm. Prepared by PL using CorelDRAW X7; www.coreldraw.com.

#### Description

The bilateral structures (Fig. 3B) consist of black, shiny amorphous carbon. The surface bears several ridges and grooves (Fig. 3A). A prominent C structure (average maximum height: 6.66 mm) with a thicker central part and thinner, proximal, curved ends is present in the center of the fossil. Opposite to the curved part, the opposing structure (“carrier“, Fig. 3A, B) comes relatively linear and only shows connections to other elements on lateral exterior. Some specimens reveal that the carriers are not necessarily fixed to the remaining structure, but more likely pass through it. The whole structure is penetrated by a widely distributed canal system whose canaliculi are calcite filled. The measured diameters of the calcite-filled (S4 Data) channels vary from approx. 7 μm (peripheral) to approx. 100 μm in the central part. The most complete specimen (NHMW 2021/0123/0057; Fig. 3B) has a maximum length of 24.24 mm, with a height of 18.94 mm. Six Type A fossils are associated with hooks, six others with a coleoid proostracum. When associated with a proostracum, this enigmatic C structure was found between 18.69 mm and 31.71 mm away from the last visible proostracum field. Generally, the distance between gladius and C structure increases with increasing proostracum length.

Occasionally, “wing-like“ structures, which sometimes appear associated with Type A fossils, are present. As the C structures apparently grow proportionally, their maxmimum height was used as indicator for the full size. The present C structures vary within a range of 3.5 to nearly 8.4 mm, depending also on low-grade diagnetic distortion. Diagenetic cracks are visible. The processus (pr in Fig. 3A, 4A) reaches lengths of 3.7–9.4 mm. The C structure to processus ratio ranges from 0.7 to nearly 1.4. Associated wing-elements can be described as triangular-elongated in shape, with length-to-base ratios from 1.5–2.2.

### Type B Fossil

Type B fossils (Fig. 3C, D and 4C, D). Black, shiny structures, triangular prolate morphology with a base consisting of three prominent interior corner points. Some specimens show an extended base. The base is slightly curved, with several ridges and grooves visible, channel system present. Large specimens reach maximum lengths of more than 10 mm (S8 Tab.). The distribution of the calcite filled channel system resembles that in Type A fossils but seems to be less developed. A prominent notch is present on the interior, but is absent on the exterior. These specimens are sometimes associated with *in-situ*-preserved coleoid hooks and with one or two additional, elongated structures. Rough surface areas were mainly found on the posterior part of the fossil. The base widths of Type B fossils were measured from 3.3–7 mm, with heights from 6.6–10.5 mm. The volumes range from 26.13–32.76 mm^3^ (S9 Tab.).

### Internal structure

Thin-sections (Fig. 5), micro probe analysis (Fig. 6) and serial sectioning confirm a prominent canal system that becomes more dense toward the center of the structures. A diameter of up to 100 μm was measured for the central canals, 7–20 μm for proximal channels. The channel system follows the geometry of the fossils. The most proximal areas of the fossils apparently lack distinct canals (at least they were not preserved).

**Figure 5.**
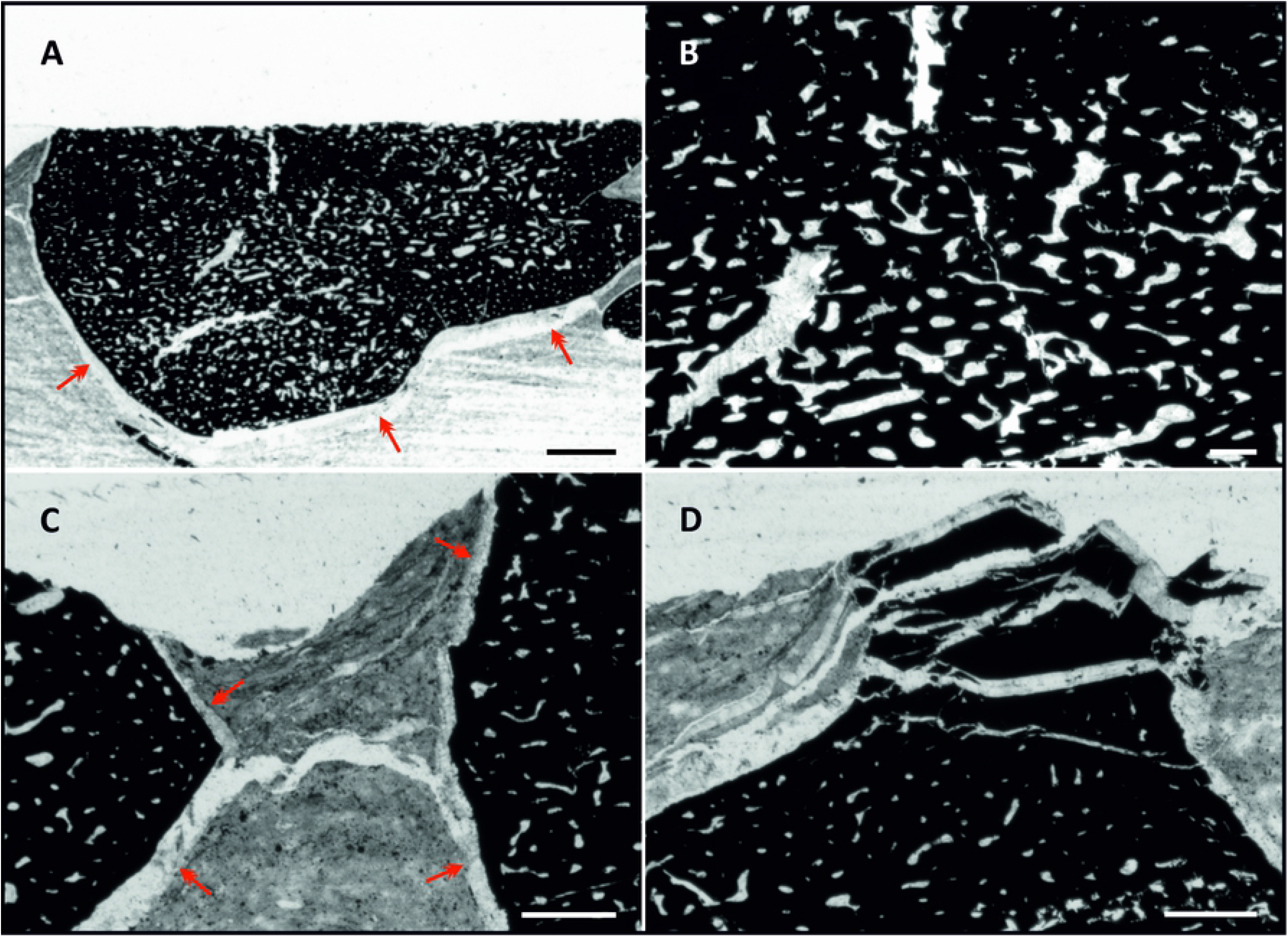
Analyses carried out on specimen NHMW 2012/0117/0024. **(A)** Thin-section of coleoid cartilage (black) reveals a widely distributed channel system (grey); red arrows: surrounding calcitic seam/layer caused by shrinking during fossilization. Note the curving of sediment around the fossil specimen. **(B)** Magnification showing pore space in the center of the fossil. **(C)** Red arrows mark the outer seam surrounding fossil show connecting areas. **(D)** Cracks in the black fossil structure caused by shrinking. All scale bars 200 μm, except scale bar for A 500 μm. Prepared by PL using CorelDRAW X7; www.coreldraw.com.

**Figure 6.**
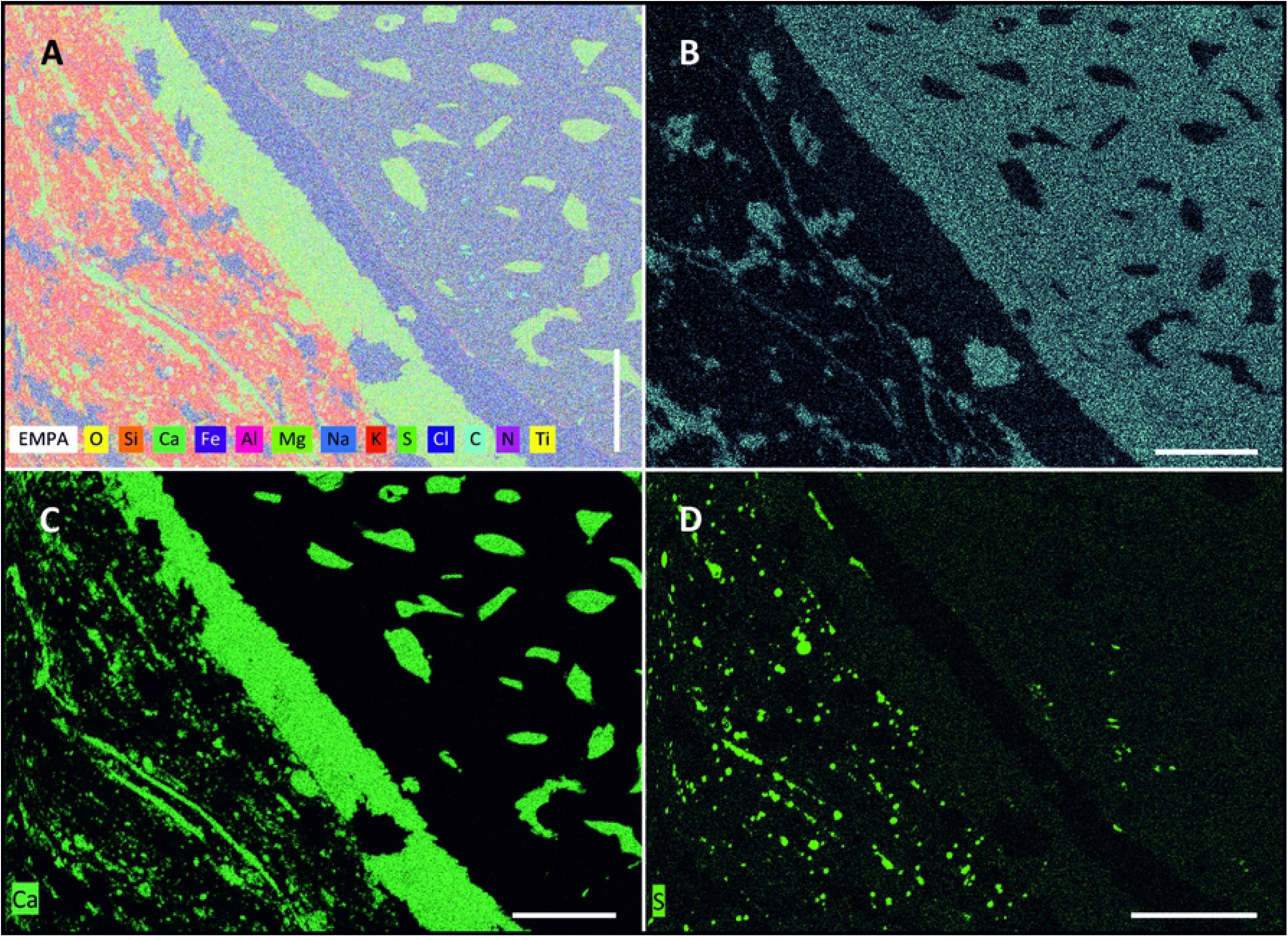
Microprobe analysis reveals the elemental distribution near the sediment-fossil boundary. A) Composite image of elements in thin-section of specimen NHMW 2012/0117/0024. Aluminium, silicium and oxygen are the main sediment constituents. **(B)** Carbon is the dominant element in the black fossil structure. **(C)** Calcite-filled channels and the fossil surrounding calcitic seam/layer. **(D)** Distribution of sulfur in the sediment corresponds to pyrite nodules. Note the outer dark layer of the fossil, lacking sulfur. All scale bars 100 μm. Prepared by PL using CorelDRAW X7; www.coreldraw.com.

### Ultrastructure and chemical composition

Microprobe elemental mapping (Fig. 6; S5 Data) of a prepared thin-section at the fossil– sediment boundary indicates a high silicium (Si), aluminium (Al), potassium (K) and magnesium (Mg) content within the sediment. Distinct nodules of sulphur (S) and iron (Fe) are visible. Carbon clearly dominates the fossil structure, but carbon-seams are also present in the sediment. Traces of titanium (Ti) are found as accumulations at the boundary of the calcitic layer surrounding the carbonised fossil. Calcium (Ca) and oxygen (O) are abundant in the fossil-surrounding layer and the channel fillings. Interestingly, S, Al and Ti limit another layer within the fossil structure. The outer surface of the fossil (approx. 50 μm) show lowest amounts of S and Ti. The transition between the outer surface of the fossil and fossil is also visible in a thin (∼ 5 μm) Al-boundary (Fig. 6A).

SEM images prove a distinct transitional layer between the carbonised fossil and calcareous sediment (Fig. 7A). The thin layer surrounding all fossils varies from 20–100 μm (Fig. 7B) and consists of the same material as the calcitic fillings (Fig. 7C, D) of the channel system, while EDS-SEM analyses confirm that the fossils consist of carbon. Raman spectroscopy of one specimen (NHMW 2021/0016/0397) (S6 Fig.) reveals a composition of strongly disordered (i.e., sp^2^ hybrid bonded) carbon for the fossil specimens.

**Figure 7.**
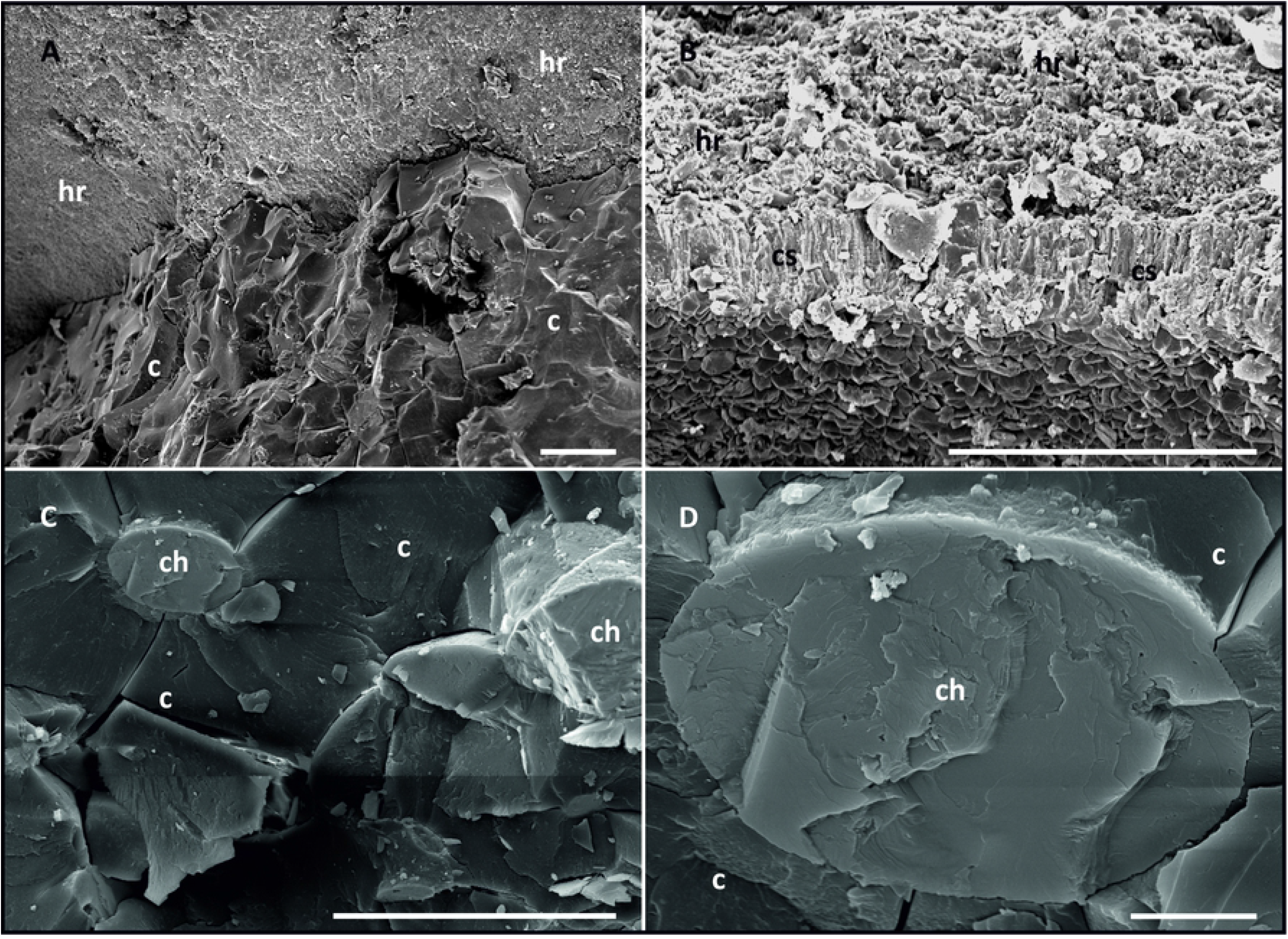
EDS-SEM images of the ultrastructure of specimen NHMW 2012/0117/0024. **(A)** Distinct boundary between fine calcareous host rock (hr) and carbonised, amorphous fossil (c). **(B)** Thin calcareous fossil-surrounding seam/layer (cs, approx. 20 μm thick). **(C)** EDX-SEM of amorphous structure with calcitic channel fillings. **(D)** Calcite-filled channel (magnification x 5000; scale bar 10 μm). hr host rock, c carbonised cartilage, ch calcitic channel fillings. Scale bar except (D) 100 μm. Prepared by PL using CorelDRAW X7; www.coreldraw.com.

## Discussion

Due to the secondary carbonisation, demineralization of the present structures in order to reinvestigate the original structure was unfeasible. Carbonised fossils are usually preserved as thin carbon-films. In contrary the here described fossils are 3D-preserved, not flattened and show no traces of abrasion or other damage. They often occur with in-situ preserved coleoid remains (phragmocone, proostracum, microhooks) from *Phragmoteuthis bisinuata*, which point to a probable coleoid origin.

Mineralization is one possibility to preserve soft tissues such as cartilage in 3D. Vertebrates require Cholecalciferol (Vitamin D) uptake for their mineralization patterns. This substance is not found in squids [50]. Recent coleoids apparently do not need cartilage mineralization in their lifecycles. Nonetheless, in vitro, several experiments showed a principal ability for mineralization of coleoid cartilage under particular circumstances [32,51,52]. Experimental in vitro coleoid cartilage mineralization requires relatively high temperatures of 37°C combined with a phosphate saturated environment [32]. Anyway it cannot be excluded that the Carnian environmental conditions (including a low-oxygen environment with high amounts of sulphur) also favoured mineralization processes after the carcasses sank to the anoxic sea floor and decay began. A previous study proposed a bacterial pseudomorphosis of coleoid soft tissues from Polzberg [12]. We agree with that idea but suggest a prior mineralization process in particular coleoid cartilaginous structures. Clearly, the morphology of the fossils differs considerably from recent coleoid head cartilage. Earlier studies indicated that the high contents of phosphatidylserine in coleoid cartilage were associated with high mineralization rates [32]. As phosphatidylserine appears to be unevenly distributed within recent coloid cartilage, this may result in incomplete mineralization rates and thus only partial preservation of cephalopod cartilage. The conspicuous, encapsulating outer calcitic layer, with abundant orthogonal cracks, is most probably caused by dehydration during carbonisation and also hints at a prior mineralization of the embedded fossils. Accumulation of pyrite crystals confirm possible euxinic conditions.

### Type A fossils

The size and morphology of the particular elements of Type A fossils closely resemble the cranial cartilage-complex (including arm cartilage, ocular cartilage and cranial cartilage with statocysts) of *Sepia officinalis* and other coleoid species (Fig. 8). In particular, the characteristic shape of the C structure resembles slightly compressed ocular coleoid cartilage (oc in Fig. 8). The ventral curvation of the C structure (Fig. 8) is also identifiable in the ocular cartilage of *Loligo vulgaris*; even the prominent groove can be matched to a corresponding groove in extant coleoids. The angle between the C structure and most probably very mobile processus is 114.4° in fossil specimens and 125.0° in recent cartilage. The preserved channel system shows a proximal–distal size-distribution, resembling the widely distributed channel system in the cranial cartilage of recent chephalopods. More channels are present in the center of the cartilage, fewer more distally (Fig. 5A). In fossils the channels with larger diameters are concentrated in the center of the structure.

**Figure 8.**
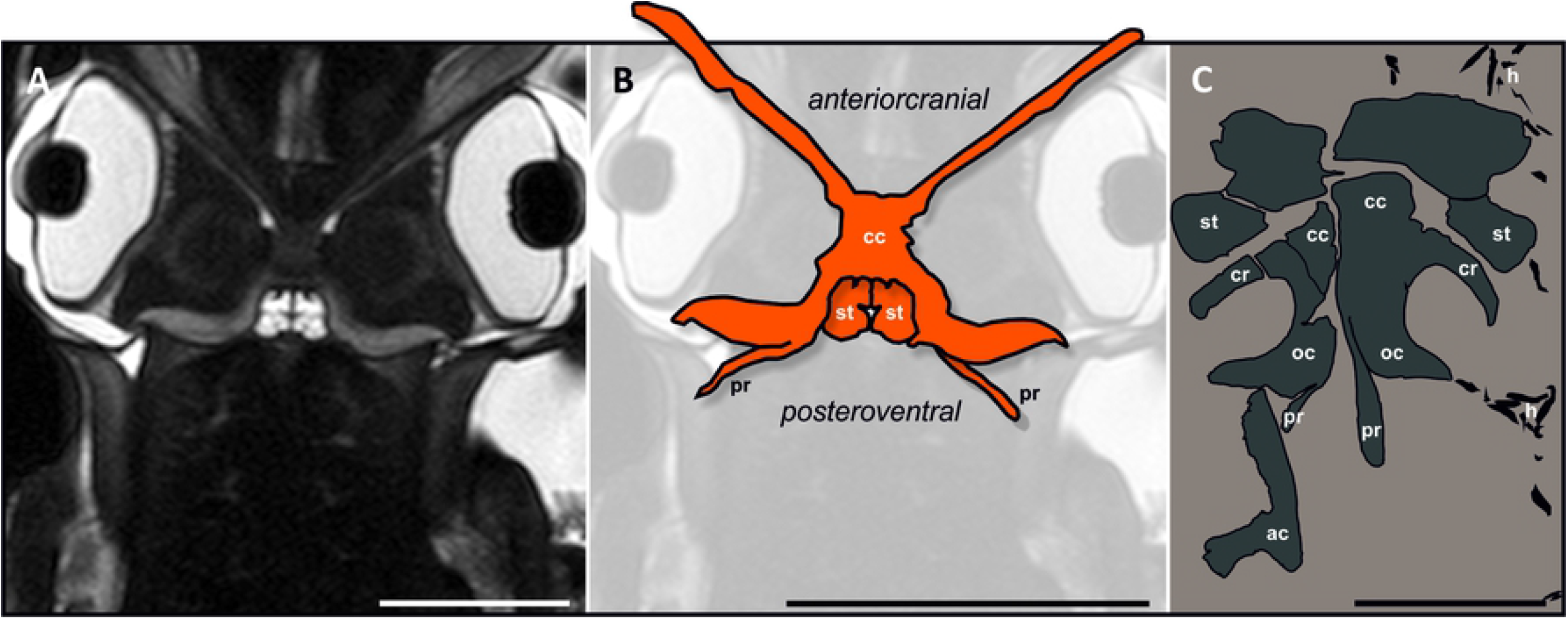
Actuopaleontological comparison of fossil specimens with recent coleoid cartilage. **(A)** MRT-data [27] of *Sepia officinalis*. Note the hook-like ending and angle to the C structure. Scale bar 20 mm. **(B)** 2D-Drawing of cartilage, redrawn magnified from a, showing principal shape of the structure in *S. officinalis*. Scale bar 2 cm. **(C)** Digital drawing of most entire specimen (NHMW 2021/0123/0057) obtained during excavations in 2021, with suggested elements as labeled: ac arm cartilage, cc cephalic cartilage, cr carrier, h hooks, oc ocular cartilage, pr processus, st statocysts. Scale bar 1 cm. Prepared by PL using CorelDRAW X7; www.coreldraw.com.

Recent, dissected specimens of *Loligo vulgaris* showed mantle lengths of 142.3–152.5 mm and head cartilage sizes of about 20 mm. Mantle lengths of dissected specimens of *S. officinalis* were 180–200 mm, with well-developed cranial cartilage lengths around 25 mm. Overall, cartilage sizes varied strongly in size, shape and thickness. Moreover, the features of head cartilage no doubt also vary with coleoid age and gender.

### Type B fossils

These mirroring specimens show conspicuous features such as the slight curvature of the prolongation to the interior and the prominent notch on the interior (Fig. 3C, D). The elements resemble the megahook-like structures of brachial crowns in Mesozoic ammonoids [53,54]. Moreover, some megahook-like structures from Polzberg show an extended “base” (which more closely resembles cartilage-elements than hooks), most probably correponding to a hook-muscular system-connective tissue. Coleoid hooks with a similar morphology have also been figured from other localites [55,56]. Belemnoids show a high variability of arm hooks (microhooks, h in Fig. 3C), also pointing to a differentation of the belemnoid arms [57]. Megahooks are geographically and taxonomically widely distributed throughout geological time [58]. Due to the introversive twist, the present structures seem well suited to fix prey or while mating. These megahooks (mh in Fig. 3C, Fig. 4C) are thought to be located at the anterio-proximal area of the coleoid [58]. Most authors propose a chitinous composition of belemnoid megahooks [58] and references therein, but a cartilaginous nature has also been discussed [59]. Although a chitinous origin is more likely, the presence of a channel system (less well-developed than in Type A fossils) points to a possible cartilaginous source material.

### Channel system

Both types of fossils have a widely branched channel system that follows the geometry of the specimens. Three thick channels were observed in the center, one of them very prominent, accompanied by smaller peripheral channels. Canal diameters of about 150 μm have been reported from canalicular fin-cartilage of an early Triassic squid-like coleoid [42]. In recent *Loligo*, the extracellular matrix is penetrated by small canals with chondrocyte extensions [18]. Original sizes of the cartilaginous channel system are about one-hundredth of the in [42] given size. Accordingly, the canals could have expanded due to the above-mentioned shrinking processes of the soft tissue during carbonisation.

### Comparison to fossil specimens from Cave del Predil (Italy)

Specimens (GBA 2006/011/0003, GBA 2006/011/0012, GBA 2006/011/0020, GBA 2006/011/0028 GBA 2006/011/0041 (Fig. 4D), all assigned to *Acanthoteuthis bisinuata* from Ladinian (Triassic) deposits of the Rinngraben ravine in Cave del Predil are known to resemble the Polzberg specimens in several ways. All the morphological elements from the Polzberg specimens could also be identified in objects from Rinngraben in Cave del Predil. Overall, the Rinngraben specimens show a poorer preservation and a more prominent appearance (see also the knob-like shape of the processus, Fig. 4D). These differences mainly reflect sedimentological and thus paleoenvironmental properties, as well as diagenetic influences. Finally, the preservation of soft tissue in *Konservat-Lagerstätten* is influenced by many interconnected factors. This complicates any clear prognosis on how they will become preserved under particular circumstances. Nonetheless, the specimens from Cave del Predil can also be interpreted as a mineralized and secondarily carbonised coleoid cranial cartilage complex.

### Actuopaleontological comparisons

The morphologies of cranial-ocular-cartilage-complexes of recent coleoid specimens such as of *Sepia officinalis* and *Loligo vulgaris* have been intensively studied. The perichondrium surrounds the whole cartilage elements and is more densely built than the remaining cartilage structure. In coronal view, the cartilage-complex in Magnetic Resonance-Tomography (MRT) data from [27] reveals a clear C structure (Fig. 7A, B), similar to the conspicuous C structure of the fossils (Fig. 7C). The presence of comparable shapes (grooves, ridges, processes, wing-like structures; Fig. 8) in ancient and recent specimens can probably be interpreted as muscular attachment sites. The well-developed canal system indicates a cartilaginous origin. The fossil specimens show peripherally thickened regions on elements of the ocular cartilage (oc in Fig. 3A, 4A), a feature also known in recent coleoids [33].

## Conclusions

Cephalopods have a long and rich evolutionary history with a high diversity of shapes and breakthrough inventions. Even the morphologies of the cephalic cartilages of the various recent cephalopod groups differ from each other in several respects. We can therefore expect a multiplicity of that morphological diversity throughout the fossil record. The exceptional conditions during deposition resulted in excellent preservation in the Polzberg *Konservat-Lagerstätte*, enabling a detailed insight into the coleoid morphology. This is the first detailed report on cranial cartilage of Mesozoic cephalopods. We analyzed the black amorphous structures with multivariate methods including thin-sectioning, SEM imaging, microprobe analyses, RAMAN and EDS measurements. All of these methods point to the cartilaginous nature of these undetermined shapes. The carbonised elements exhibit multi-tube like structures filled by secondary calcite inside the main black material. Three semi-connected elements (C structure with processus, probable brain case and statocyst capsule; connective wing-element similar in shape to the recent squids triangular pallial cartilage) form, doubled, an entire cartilage complex. Eleven analyzed specimens show mirroring pairs of elements, with smooth surfaces and protrusions serving as muscle attachments, four of them also associated with wing-elements. Additionally, numerous elements are connected with or located at the base of the basal arm hook rows, and often crowded by coleoid arm microhooks. Six specimens show the entire coleoid system with phragmocone and/or proostracum (gladius), cartilage and arm hooks, indicating and mirroring the nature and subsequently the exact position of the black carbonised elements. The described cartilage complex is direct evidence for invertebrate cartilage in Mesozoic coleoids and hence of phragmoteuthids in the Triassic ocean. The development of cartilaginous structures in marine invertebrates is still a matter of intensive discussion. Clearly, the distribution of cartilage is meaningful for taxonomic questions, whereby evidence for cartilage in fossil cephalopods can help to clarify the evolutionary role of cartilage in invertebrate groups. We present a possible solution for a 150-year-old paleontological enigma and suggest that the material represents the preservation of a mineralized and secondarily carbonised cranial-ocular-arm-cartilage complex of the gladius-bearing coleoid *Phragmoteuthis bisinuata*. Due to the secondarily carbonised preservation style of the present fossils (no demineralization possible), this interpretation is mainly based on morphological studies in fossil and extant cephalopod groups.

In a next step, the segmentation and visualisation of the canalicular system will be necessary for a conclusive explanation of the interconnected relationship between coleoid biology, cartilage mineralization processes, environmental conditions and diagenesis. Coleoid megahooks differ strongly in morphometrical properties from other known megahook occurrences. Morphometric examinations will no doubt improve our knowledge on coleoid taxonomy and ecology.

## Acknowledgements

We are grateful to Franziska and Hermann Hofreiter (Gaming), the owner of the Polzberg section for the digging permission during the whole duration of the project. We thank Birgitt and Karl Aschauer (Waidhofen an der Ybbs) for providing many fossil specimens for detailed scientific investigations. We especially thank Dr. Dan Topa (SEM, microprobe), Mr. Anton Englert (thin-sections), Mr. Goran Batic (mineralogical thin-sections) for their technical support and Prof. Dr. Lutz Nasdala for carrying out Raman Spectroscopy. Valentin Blüml and Christina Kaurin for segmentation of Micro-CT data. Leon Ploszczanski and Matthias Kranner (both Vienna) for support with SEM pictures. Martin Zuschin for dedicated supervision.

## Supporting information

**Supporting Figure S1. List of examined specimens**. The conducted methods and features on specimens such as associated coleoid remains are given. Fifty-nine specimens stem from the Polzberg locality, five from Rinngraben near Cave del Predil (Julian Alps, Italy). Indicated are inventory numbers, locality, Type A or Type B fossil and/or wing, the applied methods (Micro-CT scanning resolution is given in brackets) and further information on the specimens. Single or double element refers to the presence of only one C structure or both C structures. In specimens collected in recent excavations, the outcrop layer is given; PO Polzberg main section, ms measurements. NHMW corresponds to all inventory numbers except where GBA is given.

**Supporting Figure S2. SEM-EDS report for sample NHMW 2012/0117/0024 (Carbon)**.

**Supporting Figure S3. SEM-EDS report for sample NHMW 2012/0117/0024 (calcitic fillings)**

**Supporting Table S4. Micro-CT Scan parameters for fossil specimens**.

**Supporting Figure S5. Raman spectrum for specimen NHMW2021/0016/0397 measured at low-energy conditions**.

**Supporting Data S6. Microprobe report for sample NHMW 2012/0117/0024**.

**Supporting Figure S7. Visualisation of metrics, measured on specimens**. h_c_ height of C structure; l_p_ length of processus; m_h_ height of megahook; m_b_ base-length of megahook. Scale bars 1 cm.

**Supporting Table S8. Measurements conducted on fossil specimens**. h_c_ height of C structure; l_p_ length of processus; l_w_ length of wing; w_b_ base-length of wing element; m_h_ height of megahook; m_b_ base-length of megahook; all measurements in mm. Elements could not be measured in GBA specimens. Only specimens listed, where clear measurements were possible.

**Supplementary Table S9**. Volumes of fossil specimens, obtained from Micro-CT data.

